# 5-methoxy-N,N-dimethyltryptamine: An ego-dissolving endogenous neurochemical catalyst of creativity

**DOI:** 10.1101/578435

**Authors:** Christopher B. Germann

## Abstract

5-Methoxy-N,N-dimethyltryptamine (acronymized as 5-MeO-DMT) is *sui generis* among the numerous naturally-occurring psychoactive substances due to its unparalleled ego-dissolving effects which can culminate in a state of nondual consciousness (which is phenomenologically similar to transformative peak experiences described in various ancient contemplative traditions, e.g., Advaita Vedānta, Mahāyāna Buddhism). The enigmatic molecule is endogenous to the human brain and has profound psychological effects which are hitherto only very poorly understood due to the absence of scientifically controlled human experimental trials. Its exact neuronal receptor binding profile is a matter of ongoing scientific research, however, its remarkable psychoactivity is presumably mediated via agonism of the 5-HT2A (serotonin) receptor subtype. Anthropological/ethnopharmacological evidence indicates that various cultures utilized 5-MeO-DMT containing plants for medicinal, psychological, and spiritual purposes for millennia. In this paper we argue that this naturally occurring serotonergic compound could be fruitfully utilized as a neurochemical research tool which has the potential to significantly advance our understanding of the cognitive and neuronal processes which underpin cognition and creativity (downregulation of the default-mode network, increased neuronal functional connectivity, etc.). An eclectic interdisciplinary perspective is adopted, and we present converging evidence from a plurality of sources in support of this conjecture. Specifically, we suggest that 5-MeO-DMT has great potential in this respect due to its incommensurable capacity to completely disintegrate self-referential cognitive/neuronal processes (viz., “ego death”). The importance of unbiased systematic scientific research on naturally occurring endogenous psychoactive compounds is discussed from a Jamesian radical empiricism perspective and potential scenarios of abuse are discussed (particularly in the context of military torture).

## Introduction

The following quotation adapted from Abraham Maslow’s book “Towards a psychology of being” provides an apt primer and some semantic grounding for the subsequent essay:

> “*An essential aspect of SA* [Self-Actualized] *creativeness was a special kind of perceptiveness that is exemplified by the child in the fable who saw that the king had no clothes on - this too contradicts the notion of creativity as products. Such people can see the fresh, the raw, the concrete, the ideographic, as well as the generic, the abstract, the rubricized, the categorized and the classified. Consequently, they live far more in the real world of nature than in the verbalized world of concepts, abstractions, expectations, beliefs and stereotypes that most people confuse with the real world. This is well expressed in* [Carl] *Rogers′ phrase ″openness to experience*″ (Maslow, 1968, p. 145, content in brackets added).

Humanity is currently facing an unprecedented existential crisis which could be described as an “anthropogenic planetary emergency”. One major acute threat to the survival of the species comes from the military and the threat of nuclear annihilation, another from the destruction of the global ecosystem and the significant and extremely worrisome anthropogenic (man-made) reduction of biodiversity which will soon cause a global systemic collapse (Steffen et al., 2018). The term “biological annihilation” has been proposed to describe this ongoing scenario (Ceballos, Ehrlich, & Dirzo, 2017). Next to overpopulation (cf. Malthus), the “Anthropocene”^1^ (Lewis & Maslin, 2015) is primarily caused by the irrational, short-sighted, reckless, and ego-driven behavior of the human species, viz., overconsumption and short-term profit-oriented exploitation of the ecosystem (Fromm, 1962, 1976) – a *modus operandi* which is congruent with the philosophy of neoliberalism^2^ which is highly influential amongst the power elite (Harvey, 2007; Hill & Kumar, 2009). The continuation of the current course of action will predictably lead to total ecological catastrophe in the foreseeable future (Ceballos et al., 2017) unless humanity comes up with a radical^3^ creative solution. In 2018 the symbolic “Doomsday Clock” maintained by the *Bulletin of Atomic Scientists* has been set to “2 minutes to midnight”.^4^ According to this assessment, humanity was never that close to annihilation since 1953 when the US tested the first hydrogen bomb (Guglielmi, 2018). Therefore, creativity and a fundamentally new way of thinking are of utmost evolutionary importance if the species *Homō sapiēns sapiēns*^5^ wants to survive this century. A creative solution to this far-reaching existential problem is thus literally a matter of life or death.^6^ As Einstein put it in a New York Times interview: *“… a new type of thinking is essential if mankind is to survive and move toward higher levels*.”^7^ One of the key components of the solution is a deep understanding that the earth is a single system – an interconnected whole to which we as human beings belong.

The “single-state fallacy” (Roberts, 2006, p. 104) pertains to the widely held naïve belief that worthwhile cognition *exclusively* takes place in “normal” alert waking consciousness, a superficial assumption which fits into the contemporary materialistic and utilitarian “production-mindset” which places great emphasis on ordinary states of consciousness and disregards “altered” states of consciousness as unimportant, prejudicial, and even infantile (cf. Fromm, 1976). *Per contra*, there exists copious evidence that important creative ideas can emerge from non-ordinary states of consciousness (Tart, 1972, 2008). A well-documented illustrative historical example is August Kekulés discovery of the benzene structure in 1858, a landmark in the history of science which heralded the birth of the structural theory of organic chemistry (Kekulé, 1866, 1890). Kekulé, a German chemist, had a daydream of the *Ouroboros* (an ancient symbol of a snake seizing its own tail). This dream-image provided him with the idea of the cyclic structure of benzene (Gillis, 1966; Rocke, 2015), i.e., a symmetrical ring comprised of six carbon atoms with alternating single and double bonds. The far-reaching scientific ramifications of Kekulés insight for the rapid development of modern chemistry can hardly be overstated. Interestingly, the depth-psychologist C.G. Jung assigned specific archetypal and alchemical significance to this ancient symbol which can be found in numerous cultural traditions across various times and locations (Jung, 1969). Jung wrote: *“The dream is a little hidden door in the innermost and most secret recesses of the soul, opening into that cosmic night which was psyche long before there was any ego consciousness, and which will remain psyche no matter how far our ego-consciousness extends. For all ego-consciousness is isolated; because it separates and discriminates, it knows only particulars, and it sees only those that can be related to the ego. Its essence is limitation, even though it reach to the farthest nebulae among the stars. All consciousness separates; but in dreams we put on the likeness of that more universal, truer, more eternal man dwelling in the darkness of primordial night. There he is still the whole, and the whole is in him, indistinguishable from nature and bare of all egohood. It is from these all-uniting depths that the dream arises, be it never so childish, grotesque, and immoral.”* (Jung, 1933, p.304)

Jung’s mentor, Sigmund Freud, famously characterized dreams as “the royal road to the unconscious” (Freud, 1939). However, unbeknownst to early Freudian psychoanalysts, besides dreams, parapraxis, and free-association techniques, there are other much more effective methods to render unconscious psychic contents more accessible. Certain neuroactive chemical substances, colloquially termed psychedelics,^8^ are particularly productive tools in this regard. From a psychoanalytic perspective it is noteworthy that psychedelics produce dream-like effects and may also be classified as *oneirogenic* substances^9^ (i.e., substances that produce or enhance dream-like states of consciousness). There is a significant amount of anecdotal significant evidence that psychedelics can, *inter alia*, enhance creative ideation (indeed the term “ideagens” has been suggested; Roberts, 2006).

From a purely pragmatic vantage point on creativity the crucial importance of psychedelics in the technological development of the internet and the personal computer should be highlighted (the digital revolution). *Prima facie*, this might appear like a hyperbolic statement. However, there exists considerable historical evidence in support of the claim that psychedelics played a pivotal role in the highly creative and innovative 1960s computer-revolution which fundamentally transformed (and interconnected) the world we inhabit (see Markoff, 2005; Nelson, 1975). Besides the influence of psychedelics on the development of uniting (i.e., boundary dissolving) information-technologies like the *world-wide-web*, innumerable artists across disciplines, epochs, and cultures have been deeply inspired by transcendental experiences occasioned by psychedelics, especially within the branch termed “visionary arts” (e.g., Grey, 2001) and it has been argued unconscious processes appear to play a pivotal role in artistic expression (e.g., Kandel, 2015). Space does not permit a detailed discussion of this rich area. Eminent contemporary instances of “psychedelic-inspiration” include, for example, the entrepreneur Steve Jobs and Nobel laureate Karry Mullis.^10^ Jobs famously reported that his experience with Lysergic Acid Diethylamide (LSD) was one of the most important things he did in his whole life, a statement which recently gained experimental empirical support.^11^ Biochemist Karry Mullis was even more explicit (Mullis was honored for his ground-breaking work on the polymerase chain reaction which is today still widely used to replicate DNA). He stated in an interview: “*Back in the 1960s and early ’70s I took plenty of LSD. A lot of people were doing that in Berkeley back then. And I found it to be a mind-opening experience. It was certainly much more important than any courses I ever took*” (Schoch, 1994). He claimed that his ability to “*get down with the molecules*” was facilitated by LSD (Slattery, 2015). Moreover, he wrote in his autobiography “*The concept that there existed chemicals with the ability to transform the mind, to open up new windows of perception, fascinated me.*” (Mullis, 2000, p. 62). Mullis articulation reverberates with the title of Aldous Huxley’s classic book “The doors of perception” (Huxley, 1954) in which Huxley details his extraordinary experiences with the ancient psychedelic compound Mescaline (3,4,5-trimethoxyphenethylamine) which was administered to him by the British psychiatrist Humphrey Osmond who initially coined the term psychedelics. Huxley,^12^ a creative visionary genius who was a repeated nominee for the Nobel Prize in literature, adopted the title for his book from a phrase found in William Blake’s 1793 poem “The Marriage of Heaven and Hell”. Blake wrote: “*If the doors of perception were cleansed every thing would appear to man as it is, Infinite. For man has closed himself up, till he sees all things thro′ narrow chinks of his cavern.*”^13^ According to Huxley and Blake, overcoming the self-centered perspective associated with rigid ego-structures enables the percipient to perceive reality in fresh light and from a more impartial perspective. For obvious reasons, transcending conditioned habitual (aprioristic/automatic) perceptual schemata is crucial in the context of creative cognition. Psychedelic substances are unique in this respect because they are effective neurochemical tools which profoundly change perceptual schemata and reveal states of mind that lie far beyond the state of ordinary waking consciousness. Moreover, they possess the ability to catalyze the most “extraordinary” psychological phenomena known to science, e.g., transcendence of experiential space-time, synesthesia/somaesthesia, spectacular visual hallucinations, ineffable imaginations, intense emotional catharsis, access to unconscious/archetypal contents, profound noetic insights, enhanced biophilia, amplified empathy, etc. pp. (Gallimore, 2015; Kometer & Vollenweider, 2018; Nour, Evans, Nutt, & Carhart-Harris, 2016a). In the context at hand, one of their most important qualities is their ability to catalyze novel cognitions and perceptions and their potential to induce the process of ego-dissolution, viz., a state of non-dual consciousness^14^ (Carhart-Harris, Erritzoe, Haijen, Kaelen, & Watts, 2018; J. V. Davis, 2011; Millière, 2017; Nour et al., 2016a). In this state of nondual consciousness habitual categorical dichotomies which normally structure the all experiences are dissolved. For instance, the duality between subject and object, percipient and perceived, self and other, ingroup and outgroup, good and bad, et cetera. It has been eloquently stated that nondual consciousness is “a background awareness that precedes conceptualization and intention and that can contextualize various perceptual, affective, or cognitive contents without fragmenting the field of experience into habitual dualities” (Josipovic, 2014). The discussion of nondual consciousness has an extensive history in various ancient knowledge traditions, but it has only very recently become a research topic of neuroscience. Among experts in the field of psychedelic research, there is general consensus that psychedelics (also known as “consciousness expanding substances”) can augment cognitive processes and enable a state of “unconstrained cognition” (Carhart-Harris et al., 2012; cf. Sheldrake, McKenna, Abraham, & Abraham, 2001). Therefore, it is argued that psychedelics are important neurochemical research tools that can significantly broaden our understanding of creativity. However, this idea is not new. An early pilot study from the 1960s (which is by modern research standards unfortunately methodologically confounded) indicated that psychedelics can significantly enhance creativity and scientific problem solving (Harman, McKim, Mogar, Fadiman, & Stolaroff, 1966). After an initial phase of systematic scientific research (Carhart-Harris & Goodwin, 2017)^15^ the legal prohibition of psychedelics in the late 1960s put an abrupt halt to the short-lived research but promising agenda. The questionable “war on drugs” was initiated by the Nixon administration which had evidently dubious motives.^16^

After a long legally enforced research-hiatus, science is currently witnessing a “psychedelic renaissance”, a new rising wave of psychedelic research (Bolstridge, 2013; Cameron & Olson, 2018; Roseman, Demetriou, Wall, Nutt, & Carhart-Harris, 2018; Sessa, 2012) using modern psychological methodologies and advanced neuroimaging technologies (Carhart-Harris et al., 2012; Muthukumaraswamy et al., 2013; Roseman et al., 2016; Tagliazucchi et al., 2016). However, hitherto systematic scientific research which focuses specifically on the role of psychedelics on creativity is virtually absent and the compound 5-MeO-DMT has hitherto not been investigated in the context.^17^ We predict that future research along these lines will be very fruitful. Research on psychedelics is especially pertinent for our understanding of the neuroscience of creativity because many psychedelics have endogenous counterparts, that is, they are structurally similar or identical to neurotransmitters which constitute human physiology/neurochemistry.

Many neuroscientists are unaware that the discovery of LSD led to the idea that neurochemicals might play a role in cognitive processes. Today the fact that *neurotransmitters* influence cognition is taken for granted. However, before 1952 serotonin was thought to be a vasoconstrictor (hence the composite lexeme “sero-tonin”). In 1952-53 serotonin (5-hydroxtryptamin, 5-HT) was discovered in the brain by Betty Twarog, Irvine Page, and Sir Henry Gaddum. In 1953, Sir Gaddum took LSD in a self-experiment. Shortly afterward he and his colleague published a paper on the antagonistic effects of LSD on 5-HT (Gaddum & Hameed, 1954). Gaddum conjectured a common site of action between both compounds and theorized that the cognitive effects of LSD result from its action on 5-HT (Amin, Crawford, & Gaddum, 1954). Because he had experienced the effects of LSD first-hand he knew that it produces significant mental changes. Knowing that LSD antagonizes 5-HT, he made the novel theoretical connection for the first time in the documented history of science. That is, Gaddum was the first to postulate that 5-HT might play a role in cognition. This historical example clearly demonstrates that the systematic study of psychedelic compounds is indispensable if science wants to deepen its understanding of various cognitive processes (e.g., creativity) and their neuronal correlates. We agree with other influential creativity researchers that “evidence gleaned from the structure and function of the brain [can] enhance our ability to foster creativity” (Vartanian, 2013, p. 257; content in brackets added). Furthermore, the systematic investigation of yet uninvestigated compounds like 5-MeO-DMT might lead to novel psychopharmacological interventions and aid in the elucidation of hitherto unidentified neurotransmitter systems (cf. the discovery of the endogenous cannabinoid system). In addition, 5-MeO-DMTs molecular structure could be systematically varied (cf. A. A. Shulgin & Shulgin, 1997) in order to rigorously explore structure-activity relationships. Such research might in theory lead to the discovery of “super-agonists” (Langmead & Christopoulos, 2013).^18^ The exploration of synergistic effects with other naturally occurring psychoactive substances (e.g., Ibogaine (Glick & Maisonneuve, 1998; Winkelman, 2014)) is another hitherto uncharted and potentially very fruitful research area. In addition, allosteric modulators are of great scientific interest in this context (cf. Schwartz & Holst, 2007). That is the agonistic actions of 5-MeO-DMT can in principle be enhanced (>100% efficacy) by various allosteric modulators (e.g., via allosteric modulators of G protein-coupled receptors; cf. May, Leach, Sexton, & Christopoulos, 2007). Yet another related important research question concerns the “entourage effect” (cf. Sanchez-Ramos, 2015). That is, what are the neuropsychopharmacological and phenomenological differences between the pure compound (synthesised 5-MeO-DMT) and the compound as found in nature within a whole complex organism (toad venom, tree bark, seed pods, etc.)?

In order to provide corroborating empirical evidence that psychedelics are important research tools in the context of creativity research we will now discuss two more recent experimental studies which are pertinent to the psychology and neuroscience of creativity. Based on the relevant literature (e.g., Nour, Evans, Nutt, & Carhart-Harris, 2016), we specifically argue that an understanding of the psychological and neurophysiological processes which undergird ego-dissolution is pivotal for advancing our scientific understanding of creativity. After introducing the corroborating studies, we will provide more detailed information on the underappreciated and virtually unresearched endogenously occurring compound 5-MeO-DMT. Based on this background we will then formulate several empirically falsifiable hypotheses.

## Psilocybin increases “Openness to Experience”

Psilocybin (O-phosphoryl-4-hydroxy-N,N-dimethyltryptamine) is an indole alkaloid (a structural relative of 5-MeO-DMT)^19^ which was synthesized and named by the Swiss chemist Albert Hofmann^20^ (Hofmann et al., 1959; 1958). The compound is present in more than 150 fungi species, some of which are endemic to the USA and Europe (e.g., *Psilocybe semilanceata*, known as Liberty Cap). In shamanic contexts, psilocybin has been utilized for spiritual and healing purposes for millennia.^21^ Its molecular structure closely resembles 5-hydroxtryptamine (5-HT, serotonin). In humans, psilocybin functions as a prodrug and is rapidly dephosphorylated to psilocin (4-N,N-dimethyltryptamine) which acts as a non-selective partial 5-HT receptor agonist. It shows particularly high binding affinity for the 5-HT1A, 5-HT2A, and 5-HT2C receptor subtypes (Kraehenmann et al., 2015; Nichols, 2004). A landmark study conducted at Johns Hopkins University by MacLean, Johnson, & Griffiths (2011) experimentally demonstrated that a single high-dose of psilocybin can induce long-lasting personality changes in the domain “Openness to Experience”, as measured by the widely used NEO-PI (Personality Inventory).

Openness to Experience (OTE) is one of the core dimensions of the extensively employed quinquepartite (big five) model of personality. OTE is an amalgamation of several interconnected personality traits which include: 1) aesthetic appreciation and sensitivity, 2) fantasy and imagination, 3) awareness of feelings in self and others, and 5) intellectual engagement. Most relevant for the context at hand is the fact that OTE has a strong and reliable correlation with creativity (Ivcevic & Brackett, 2015; S. B. Kaufman et al., 2016; Silvia, Nusbaum, Berg, Martin, & O’Connor, 2009)^22^. Individuals with high scores on the OTE dimension are “permeable to new ideas and experiences” and “motivated to enlarge their experience into novel territory” (DeYoung, Peterson, & Higgins, 2005). The experimentally induced increase in OTE was mediated by the intensity of the mystical experience occasioned by psilocybin. Importantly, ego-dissolution is a central feature of mystical experiences (see also Griffiths, Richards, McCann, & Jesse, 2006). Hence, it is logically cogent to assume that the experience of ego-dissolution correlates significantly with increases in openness to experience.

## LSD expands global connectivity in the brain

A recent multimodal fMRI study by Tagliazucchi et al. (2016) conducted at Imperial College London administered LSD-25 intravenously to healthy volunteers. The researchers found that LSD-induced ego-dissolution was statistically significantly correlated with an increase in global functional connectivity density (FCD) between various brain networks (as measured by fMRI) indicating that the psychedelic enabled novel configurations of brain states. As discussed in the previous study by MacLean et al. (2011), mystical experience (ego-dissolution) is correlated with an increase in OTE which in turn is strongly correlated with creativity.^23^. One of the key findings of the fMRI-study was that high-level cortical regions and the thalamus displayed increased connectivity under the acute influence of LSD. To be specific, increased global activity was observed bilaterally in the high-level association cortices and the thalamus (often regarded as the brains “central information hub” which relays information between various subcortical areas and the cerebral cortices). The global activity increase in the higher-level areas partially overlapped with the default-mode, salience, and frontoparietal attention networks. The FCD changes in the default-mode and salience network were predicted a priori due their association with self-consciousness. As predicted, a significant correlation between subjectively reported ego-dissolution and increase in global connectivity between networks was detected. The results of this important study demonstrate for the first time that LSD increases global inter-module connectivity while at the same time decreasing the integrity of individual modules. Specifically, LSD enhanced the connectivity between normally separated brain networks (as quantified by the widely used Φ connectivity/associativity index^24^). The observed changes in activity significantly correlated with the anatomical distribution of 5-HT2A receptors. These findingst is especially relevant for researchers who want to identify the neural correlates of creativity because an enhanced communication between previously disconnected neuronal network modules is assumed to be crucial for the generation of novel ideas (e.g., Moore et al., 2009). Moreover, associative processes are generally assumed to play a key role in creativity (Lee, Huggins, & Therriault, 2014). Tagliazucchi et al. concluded that LSD reorganizes the rich-club architecture of brain networks and that this restructuring is accompanied by a shift of the boundaries between self and environment. That is, the ego-based dichotomy (i.e., dualism) between self and other, subject and object, internal and external, dissolves as a function of specific connectivity changes in the modular networks of the brain^25^. Taken together, Tagliazucchi et al. (2016) demonstrate that LSD induced ego-dissolution is accompanied by significant changes in the neuronal rich-club architecture and that ego-dissolution is accompanied by the downregulation of the default-mode network (DMN).^26^ In the context of creativity research this finding is particularly intriguing because the DMN is associated with habitual thought and behavior patterns which are hypothesized to be negatively correlated with creativity and the generation of novel ideas. That is, downregulation of the DMN by psychedelics and the accompanying phenomenology of ego-dissolution are promising factors for the understanding (and enhancement) of creativity (cf. Vartanian, 2013). Based on these findings, we suggest a novel neuropsychopharmacological mechanism for the enhancement of creativity which has, to our best knowledge, never been proposed before. Our hypothesis highlights the importance of ego-dissolution for the enhancement of creativity. That is, a reduction of the influence of the self-referential ego structures (presumably mediated via DMN disintegration) on perception and cognition enables the perception of reality from a new (more unbiased/impartial) perspective, thereby enabling perspectival multiplicity and cognitive flexibility which is crucial for creative ideation. Based on the conjecture that ego-dissolution provides a “cognitive reset” which enables humans to perceive and conceptualize reality from a more unconstrained non-dualistic perspective, we argue that 5-MeO-DMT is an especially intriguing molecule in this regard because its ego-dissolving effects are much more pronounced than those of psilocybin or LSD (or any other known psychedelic). The “reset theory” is a first attempt to formulate a causal mechanism which could explain why ego dissolution associated with the hypothesized increase in creativity. Ego-dissolution could enable humans to “See things with new eyes” — i.e., via a reduction of the structuring and organizing influence of perceptual schemata and a reduction of inhibitory top-down influences^27^ (i.e., preconception vs. apperception). Empirical data indicates that ego-dissolution is a unique property of psychedelic substances (Nour et al., 2016a). In a web-based study utilizing the Ego-Dissolution Inventory (EDI) several substances were compared and the results showed that only psychedelics were significantly correlated with the experience of ego dissolution. *Per contra*, other psychoactive substance like alcohol or cocaine enhance an egoic style of cognition (ego inflation).^28^. In the same study, participants also responded to a subset of items from the Mystical Experiences Questionnaire (MEQ) (Barrett, Johnson, & Griffiths, 2015). The results indicated a positive correlation between psychedelic dose and the strength of the mystical experience. As discussed, a defining feature of the mystical experience is a ego-dissolving “unitive” (nondual) experience.

This has already been pointed out by William James more than a century ago (James, 1985/1902). Unity experience is closely related to the Freudian concept of “oceanic feeling” (oceanic boundlessness) — a sensation of being one with universe. In fact, Romain Rolland formulated the phrase in a letter to Freud. Rolland argued that it is this nondual experience which lies at the core of all religious feelings (theistic or nontheistic). Freud utilized this idea in his later writings and hypothesized that this nondual state of consciousness is a psychological residue from the infantile stage in which the egoic schism between self and other (object and subject) has not yet occurred (Freud, 1930). That is, according to Freud nondual experiences are a relict of the developmental stage in which the newborns formation of the self-concept has not yet taken place and has consequently not yet divided experience (perception) into permanent self versus non-self dichotomies.

## 5-MeO-DMT: An endogenous catalyst for creativity

The intranasal administration of 5-MeO-DMT in form of a snuff preparation called “Cohoba”^29^ by the Taíno people of Hispaniola was first observed around 1496 by Friar Ramón Pané who reported his observation to Christopher Columbus who in 1492 made initial contact with this culture (Nunn & Qian, 2010; Shultes, 1976; Torres & Repke, 2006). In the context of contemporary science, 5-MeO-DMT is a relatively unknown member of a group of naturally-occurring psychoactive indolealkylamines (Glennon & Rosecrans, 1982; A. T. Shulgin & Carter, 1980). It was first synthesized by Japanese chemists in 1936 who published their results in German (Hoshino & Shimodaira, 1936). The tryptamine is an analog of tryptophan and endogenous to human physiology. Research indicates that 5-MeO-DMT may be endogenously synthesized in human pineal and retina. Moreover, it has been detected in blood, urine, and cerebrospinal fluid (Shen, Jiang, Winter, & Yu, 2010). Its extremely powerful acute effects are pharmacokinetically short-lived. As many other tryptamine psychedelics, it acts as a nonselective 5-HT agonist and causes a broad spectrum of highly interesting psychological, effects. It displays a high binding affinity for the 5-HT1A and 5-HT2 and subtypes (Krebs-Thomson, Ruiz, Masten, Buell, & Geyer, 2006) but other mechanism of actions appear to be involved in its psychoactivity (e.g., inhibition of enzymatic monoamine oxidase activity; but see Nagai, Nonaka, & Satoh Hisashi Kamimura, 2007). The 5-HT system is associated with, cognition, emotion, and memory, *inter alia*. For example, 5-HT receptors are located in the cerebral cortex (cognition), in the amygdala (emotions), and in the raphe nucleus (its projection regulate circadian rhythms, alertness, inhibition of pain, *inter alia*). The raphe nucleus is located in the phylogenetically most primitive part of the brain, the brainstem, and its serotonergic axons project widely throughout the cortex. The raphe nucleus produces the majority of brain serotonin and it contains ≈85% of all the of the serotonin neurons in the brain (Hornung, 2003). Ergo, when it is stimulated by 5-MeO-DMT it causes extensive serotonergic activation throughout many interconnected neural networks of the brain. Moreover, 5-HT receptors are present in the hypothalamus which connects the central nervous system to the endocrine system. It can be cogently argued that 5-MeO-DMT causes the hypothalamus to release significant amounts of the neuropeptide oxytocin via the pituitary gland. This hypothetical increase in oxytocinergic activity might explain why the qualitative linguistic descriptions of 5-MeO-DMTs phenomenology frequently include words like “love”, “unity”, and “connectedness”. Accumulating evidence indicates that 5-MeO-DMT is an endogenous ligand of the Trace amine-associated receptors (TAARs), a class of G protein-coupled receptors that were only recently discovered in 2001 (Carbonaro & Gatch, 2016). It has been hypothesized that TAARs are involved in sensory perception (Wallach, 2009). Moreover, TAARs have been associated with pathological neuroadaptations associated with prolonged exposure to addictive drugs (e.g., alcohol, heroin, cocaine, etc.). Consequently, this molecular target might partially explain 5-MeO-DMTs promising neurorestorative and neuroprotective effects (Dakic, 2017). Because 5-MeO-DMT is able to target these receptors it might be able to regulate the pathological neurological adaptions, for example those caused by various substance (and possibly behavioural) addictions (cf. the neuropsychological “reset-hypothesis” (cf. Carhart-Harris et al., 2017)). Hence 5-MeO-DMT might counteract rigid cognitive and behavioural patterns, thereby facilitating cognitive flexibility (cf. Gruner & Pittenger, 2017). In support of this view, a recent cutting-edge *in vivo* and *in silico* study using human cerebral organoids (Dakic et al., 2017) demonstrated that 5-MeO-DMT has modulatory effects on neuroplastic processes, long-term potentiation, cytoskeletal reorganization, and microtubule dynamics (cf. Hameroff & Penrose, 2014). Specifically, it was found that 5-MeO-matches the σ1 receptor which regulates cytoskeletal dendritic spine morphology and neurite outgrowth (Dakic et al., 2017). Therefore, σ1 receptor agonism may mediate neuroplastic processes which are crucial for creativity, cognitive flexibility, and sustained cognitive/behavioural changes (Sun et al., 2016). In addition, agonism of the σ1 receptor has been shown to have anti-inflammatory effects (Szabo, 2015) which may positively influence various creativity related cognitive processes.

5-MeO-DMT is widespread in the plant kingdom and has been used in shamanic rituals for millennia (Torres et al., 1991). While its structural relative Psilocybin is exclusively present in numerous fungi species, 5-MeO-DMT is present in various plants, for instance *Virola theiodora* (Agurell et al., 1969), a tree species belonging to the *Myristicacea*e (nutmeg) family. In additions to its relatively widespread phytochemical distribution, it is present in high concentrations in the venom of *Incilius alvarius* (known as the Sonoran Desert toad), an *Amphibia* which produces significant amounts of 5-Meo-DMT in its numerous parotoid glands as a defensive chemical mechanism against predators (Erspamer, Vitali, Roseghini, & Cei, 1965; Hutchinson & Savitzky, 2004). The salience of toad symbolism in Mesoamerican art and mythology is remarkable and well documented by anthropologists and toad effigies (with oftentimes accentuated glands) are prominent in the historical remains of the Mayan and Aztec cultures (Davis & Weil, 1992).^30^ Moreover, 5-MeO-DMT can sometimes be found in certain variations of Ayahuasca (a drinkable plant-based concoction, which is utilized by indigenous tribes in the Amazonian rainforest for divinatory and healing purposes), for instance, when the leaves of the plant “Chaliponga” (*Diplopterys cabrerana*) are added to the concoction (Callaway et al., 2006; Rätsch, 1998). 5-MeO-DMT has been utilized for spiritual purposes as a religious sacrament in the rituals of the USA based Christian “Church of the Tree of Life” (Gottlieb, 1994). Modern artworks inspired by 5-MeO-DMT experiences are oftentimes geometrically highly complex and depict multidimensional fractal-like symmetric mathematical structures^31^ an observation which is particularly intriguing from a neuroesthetics point of view (cf. Ramachandran & Hirstein, 1999). Despite its longstanding usage in the course of human evolution^32^, systematic research human trials are currently lacking, and science does not know much about the psychological effects of 5-MeO-DMT. The research area is thus truly uncharted novel scientific territory (and its exploration required openness to experience on the part of the research community; ibid., p. 13). It has been convincingly argued that it is of is of “potential interest for schizophrenia research owing to its hallucinogenic properties” and that research on 5-MeO-DMT can “help to understand the neurobiological basis of hallucinations” (Riga, Soria, Tudela, Artigas, & Celada, 2014)^33^ even though visual hallucinations are much less commonly reported compared to its structural analog N,N-Dimethyltryptamine (DMT) which induced the most spectacular vivid visual perception possibly imaginable (but see Strassman, 2001).

5-MeO-DMT exerts extremely profound effects on the self-concept (ego). Here, the term ego is not used as defined in the classical Freudian tripartite model (Freud, 1923), but it refers to the concept of encapsulated identity i.e., who we think we are as human beings. Thus, the usage of the term ego is more closely aligned with the ancient Sanskrit term “Ahaṃkāra” as defined in Vedic philosophy (cf. Cartesian positional identity; Comfort, 1979). In this theoretical/phenomenological framework, the ego can be conceptualized as a filter or a lens which converts experiences. Pure consciousness, on the other hand, lies beyond the ego construct and is “that which perceives” (cf. Josipovic, 2010, 2014). While the ego identifies with the content of sense experience, consciousness itself does not (Sivananda, 1972). Coonsciousness itself has no associated identity. It is a detached witness of experience.^34^. 5-MeO-DMT by far the most effective pharmacological agent for ego-dissolution and it is much more potent than its structural relatives (e.g., N,N-Dimethyltryptamine), qualitatively and quantitatively. It has been described as a prototypical entheogen (Metzner, 2015).

An entheogen (Ruck, Bigwood, Staples, Ott, & Wasson, 1979) is a chemical substance used in a religious, shamanic, or spiritual contexts that has the potential to produce profound psycho-spiritual insights and changes. The etymology of the neologism “entheogen” is a compound lexeme derived from the ancient Greek ἔνθϵος (entheos) and γϵνέσθαι (genesthai) and translates into “generating the divine within” (cf. “enthusiasm”). 5-MeO-DMT is a ceremonial sacrament (Eucharist) of the “Church of the Tree of Life” and interdisciplinary research focusing on 5-MeO-DMT might provide further impetus for the emerging new neuroscientific paradigm which goes by the name “neurotheology” (Winkelman, 2004). Following this line of thought it has been stated by the eminent neurobiologist Efrain C. Azmitia that “*the ability of these drugs to induce a feeling of closeness to God is a special property of the indoles and this property is attributed to activation of the cortical 2A serotonin receptor*” (Azmitia, 2012).

We would like to repeat the crux of our argument: Given its phenomenological profundity and its unparalleled efficiency to dissolve ego structures we propose that 5-MeO-DMT should be systematically investigated in order to elucidate the postulated connection between non-dual (ego-less) states of consciousness and the associated enhancement in creativity. One pillar of this hypothesis is the idea that ego-dissolution is associated with a breakdown of linguistic structures^35^ (hence the characteristic ineffability of its phenomenology). According to the Saphir-Whorf hypothesis of linguistic relativism, language structures cognition and perception in significant ways. Ergo, we hypothesize that a release from the strong aprioristic influences of linguistic processes enables a more unrestrained style of cognition and perception. Further, we argue that the collapse of the “subject versus object” dichotomy into a non-dual experience has enormous potential for complex cognitive restructuring (cf. Josipovic, 2010). “Ego death” (ego-dissolution) is emotionally and cognitively extremely challenging which resonates with the “hardship model of creativity” (Forgeard, 2013). At the same time the extremely challenging experience of ego-dissolution may also have significant positive therapeutic/cathartic effects which are of importance in the context of creativity research (e.g., release from severe trauma, stress, unconscious tensions). The experiences induced by 5-MeO-DMT are tremendously radical^36^ and therefore capable to disperse deeply engrained cognitive/perceptual schemata^37^, thereby enabling a more unrestricted style of cognition^38^. Specifically, we argue that due to its unparalleled ego-dissolving properties 5-MeO-DMT facilitates a less self-centered and hence more unbiased style of cognition which enhances creativity. This hypothesis is empirically falsifiable in the Popperian sense and various established cognitive testing procedures^39^ could be utilized to test this hypothesis experimentally. The logical which undergirds our theorizing can be formalized using propositional logic, i.e., in form of a deductive syllogistic argument.

Major premise: Ego-dissolution enhances creativity.

Minor premise: 5-MeO-DMT induces ego-dissolution.

Deductive conclusion: ∴Ergo, 5-MeO-DMT enhances creativity.

Based on this argument, we postulate the following directional hypotheses:

**H_1_:** Self-reported ego-dissolution predicts subsequent enhancements in creativity, as quantified by various creativity test batteries (e.g., J. C. Kaufman, 2012) in a dose-dependent manner. This effect is mediated by the profundity of the experience, e.g., how challenging the experience was, intensity of the “peak experience”, personal meaningfulness of the experience, etc. (Barrett, Bradstreet, Leoutsakos, Johnson, & Griffiths, 2016; Forgeard, 2013; Griffiths et al., 2006; Majić, Schmidt, & Gallinat, 2015).
**H_2_:**The intensity of 5-MeO-DMT induced ego-dissolution predicts consequent increases in aesthetic perception, biophilia, and feelings of fundamental existential interconnectedness^40^ (viz., “nonduality”).
**H_3_:**The intensity of ego-dissolution experienced by participants predicts the longitudinally measured significance of the life-event in a non-linear dose-dependent manner, similar to the pattern observed in studies with the structural analog psilocybin (Griffiths, Richards, Johnson, McCann, & Jesse, 2008).

*Ex hypothesi*, we argue that the conjectured effects are objectively quantifiable and reliably replicable in a rigorously controlled experimental setting. Up to date, we are unaware of any systematic scientific research which focused specifically on the effects of 5-MeO-DMT on ego-dissolution and creativity. Consequently, we suggest that future studies should be designed in order to elucidate this rich and potentially very fruitful research area. The present discussion is just a start in order to motivate future studies along these lines.

## Brains in chains: Neuropolitics, neurodiversity, and cognitive liberty

The UK is the first country in human history which generically bans all psychoactive substances, i.e., an omnibus prohibition of all mind-altering chemicals, irrespective of their well-documented historical use and their safety profile, for example, as objectively quantified by the conventional LD_50_ and TD_50_ toxicity indices. For instance, psilocybin exhibits remarkably low toxicity. The LD_50_ in humans remains unknown, given the lack of any intentional or accidental poisoning death data. The therapeutic window (or pharmaceutical window) is extremely safe and the maximum tolerated dose (MTD) is very high, i.e., the therapeutic index is very high.^41^ A common metric in comparative risk assessment is the margin of exposure (MOE), defined as the ratio between the toxicological threshold (defined as the benchmark dose) and the estimated average human intake (Lachenmeier & Rehm, 2015). Both, MTD and MOE indicate a very benign safety profile for psilocybin, especially compared to neurotoxic agents like alcohol which has a very low MOE (Lachenmeier & Rehm, 2015) and has been associated with numerous detrimental neurocognitive (Weitemier & Ryabinin, 2003), genetic, and epigenetic effects (Chen, Ozturk, & Zhou, 2013). Despite these facts, psilocybin is classified as a class A substance in the UK. This Regulation is a *lex specialis*, which introduces an additional serious burden for researchers interested in non-ordinary states of consciousness. Regrettably, scientific research on psychedelics is currently legally highly restricted due to the irrational Class A status of psychedelic substances as defined in the “Psychoactive Substances Act^42^” (PSA) which reached Royal Assent in January 2016. The PSA generically prohibits *all* mind-altering substances besides the most harmful and addictive ones which are of commercial significance (e.g., alcohol and tobacco; but see Nutt, King, & Phillips, 2010) and it classifies relatively harmless substances like psilocybin on par the most harmful and detrimental substances like heroin, cocaine, and alcohol. This classification is based on the assumption that psilocybin has no medicinal value which is clearly not the case (but see Bogenschutz & Johnson, 2016). The widespread psychological propaganda (Bernays, 1928, 1936; Mullen, 2010) against psychedelics (linking psychedelic use to psychopathology and suicide) which was initiated by Nixon’s “war on drugs” campaign, has now been evidently debunked (Johansen & Krebs, 2015), even though the public mind is still under its influence. Well informed legal scholars interpret the PSA as an explicit violation of the right to mental self-determination (i.e., cognitive liberty; Walsh, 2016) - particularly in the context of Article 9 of the European Convention on Human Rights which should protect the right to freedom of thought. It is obvious that cognitive liberty is a prerequisite for creativity. It can be convincingly argued that the PSA reduces neurodiversity - it homogenizes^43^ cognitive/neuronal processes and restricts memetic and, ergo, cultural evolution (in analogy with the importance of genetic diversity in the context of biological evolution). *Summa summarum*, the PSA is not evidence-based and presents a serious legal impediment to scientific research, creativity, and cognitive innovation (see also Boire, 2000).

## Potential for military abuse of psychoactive substances

Given the fact that 5-MeO-DMT is unparallel in its ability to almost instantaneously dissolve psychological ego-structures it can in principle be utilized as “psychochemical weapon”, for instance, in the context of military operations. In the past, methods that enable the dissolution of ego structures have been of great interest to the military in the context of interrogation and torture (e.g., the work of Donald Hebb on sensory deprivation).^44^ Only recently several APA psychologists were associated with psychological torture in the context of military operations which led to a refusal of the APA to participate in future operations.^45^ 5-Meo-DMT can be extremely destructive to the human psyche when utilized for the wrong purpose and in the wrong setting. Therefore, it is of utmost importance to develop legal frameworks which prevent its application in “situations of crisis” (so called ticking-bomb scenarios) in which the principles laid down by the human rights convention (e.g., Geneva Conventions and the U.N. Convention Against Torture) are thought to be no longer applicable for utilitarian juridical reasons (cf. Horowitz). Given the rapid breakdown of 5-MeO-DMT within the human system (pharmacokinetic elimination) it is in principle difficult to prove its illegal application post hoc. The application of 5-MeO-DMT in a military context can have disastrous consequences because the compound *completely* breaks down psychological defense mechanisms. A person under the influence of 5-MeO-DMT is utterly defenseless and the “interrogator” has consequently “god-like sovereignty” (cf. Améry, 1966) to subject the individual to all kinds of psychological interventions which are otherwise impossible. Furthermore, stimuli which are normally perceived as relatively harmless will be perceived as extremely threatening and their impact will be amplified in unpredictable ways – thereby causing irreversible psychological damage. Physical torture is always also psychological torture, but it leaves the possibility to distance (dissociate) psychologically from the torturer which allows in principle for a partial coping with the traumata. Psychological torture, on the other hand, targets the very core of a human being and therefore destroys the entire person and not just his/her physical body. In the hands of the wrong people 5-MeO-DMT is an extremely cruel and destructive psychological weapon which can induce a form of permanent trauma which is unimaginable to a normal person as it intervenes into the core dynamics of consciousness and allows for radical modifications of the ego. For obvious reasons, it is a two-sided (neurochemical) sword which can be used to foster human development or to manipulate and *deeply* traumatize human beings. It follows on ethical/moral grounds that an absolute (deontological) categorical prohibition of the use of 5-MeO-DMT for military/political purposes is of great importance, especially in the present historical context in which basic human rights have been repeatedly violated under dubious political motives (e.g., within the justificatory/exculpatory utilitarian frame of “national security”). We cannot allow for double standards and moral elasticity when it comes to human consciousness itself! (cf. Immanuel Kant’s foundational work on morality)

## Conclusions

We would like to close by reconnecting the topic back to the introduction of this essay. In a recent PNAS article entitled “Trajectories of the Earth System in the Anthropocene” it has been stated that: “***Collective human action** is required to steer the Earth System away from a potential threshold and stabilize it in a habitable interglacial-like state. Such action entails stewardship of the entire Earth System-biosphere, climate, and societies-and could include decarbonization of the global economy, enhancement of biosphere carbon sinks*, ***behavioral changes***, *technological innovations, new governance arrangements, and **transformed social values***” (Steffen et al., 2018, p. 8252; emphasis added by the author).

We strongly agree with this conclusion (note that “Earth System” is used in the singular not in the plural). Given the “extraordinary danger of the current moment” (see Doomdayclock statement, 2018)^46^ it undeniable that we as human beings need to radically change our egoistic behavior as a species otherwise our existence on this planet will come to a catastrophic end soon. Behavior is governed by thought and the basis of thought is consciousness. Consequently, the essential question is: How can human consciousness can be transformed for the better to change the trajectories of the Earth System? Science (and particularly neuroscience and psychology) play a central role in answering this question and the neurochemical correlates of consciousness play a pivotal role in this context. So far, contemporary science has largely neglected the extraordinary experiences catalyzed by psychedelics and the potential of the powerful endogenous neurochemical 5-MeO-DMT has not yet been explored at all. Specifically, research in the domain of creativity appears to be potentially very fruitful and given the pressing urgency of the situation 5-MeO-DMT should be systematically investigated as soon as possible. If there is a chance that chemicals like 5-MeO-DMT can catalyze a radical new (less egocentric) way of thinking which fosters biophilia and personal insight into the interconnectedness of nature and all human beings, then it is sciences moral obligation to take this potential very seriously as creative change is a matter of survival. The transformational ego-dissolving experience of nonduality might prove be the quintessential antidote to the rigid habitual, materialistic, dualistic, and egoic mindset which lies at the very core of the existential crisis humanity is facing. That is, an egoic mindset is incompatible with the urgent need for collective action. Specifically, the nondual experiences occasioned by psychedelic compounds like 5-MeO-DMT appear to be antagonistic towards the dualistic egocentric paradigm (a narcissistic consumer-mindset) which regards nature as an exploitable resource. In other words, the widely shared and culturally conditioned dualistic psychological perspective which separates man from nature stands in sharp contrast with the interconnected unitive worldview catalyzed by this extraordinary ego-dissolving tryptamine (Carhart-Harris et al., 2018; Lyons & Carhart-Harris, 2018a; Nour, Evans, & Carhart-Harris, 2017). Furthermore, epistemological insights into the nondual ontology of existence (e.g., dual-aspect monism/neutral monism)^47^ challenge the core assumptions of contemporary science, viz., the notion of detached objectivity which is *de facto* a cognitive illusion (Hoffman, 2016; cf. Wiseman, 2015). A nondual conceptualization of reality might force us to rethink our most fundamental (but non-evidence based and naïve) beliefs about the way we conceive reality and practice science, e.g., the stipulated dichotomy between observer and observed^48^ and the widespread belief that the brain *produces* consciousness.^49^ A nondual reconceptualization is therefore implicitly perceived as a threat to the widely adopted “quasi-Newtonian” *status quo*^50^ which has in reality already been fundamentally revised by modern quantum physics (viz., the widely held and mainly unquestioned metaphysical assumption of local realism is no longer tenable (Wiseman, 2015)). However, most of science still operates under an outdated deterministic Newtonian paradigm. In his classic book “The structure of scientific revolutions” Thomas Kuhn pointed out that it is general phenomenon that paradigm challenging anomalies “that subvert the existing tradition of scientific practice” (Kuhn, 1970, p. 6) are neglected as long as possible. Along the same line, Abraham Maslow discusses the “Psychology of Science” in great detail in his eponymous book (Maslow, 1962). Maslow formulates a quasi-Gödelian critique of orthodox science and its “unproved articles of faith, and taken-for-granted definitions, axioms, and concepts”. Research on extremely powerful consciousness-altering substances like 5-MeO-DMT might force us the rethink our most fundamental beliefs about the way we conceive reality and practice science. *Prima facie*, this line of thought might sound absurd to the majority of critical readers. However:

> “*If at first the idea is not absurd, then there is no hope for it*.”

~ Albert Einstein (as cited in Hermanns & Einstein, 1983)

Moreover, it might be helpful to look at the way indigenous cultures which utilised 5-MeO-DMT related to the earth (and to each other). It might be argued that ego-dissolving psychoactive plants and fungi played an important role in this relationship. Of course, such a solution sounds absurd to the modern mind. However, any real solution to the “anthropogenic global crisis” will be at odds with the predominant *status quo* and will thus cause intense cognitive dissonance. If science wants to live up to its ideal to capture reality in its entirety without leaving any residue it needs to integrate neurochemicals like 5-MeO-DMT into its modelling efforts - especially given the fact that this alkaloid is an endogenous components of the human brain and ergo arguably of evolutionary relevance. Any model which incorporates only a specific (*a priori* selected) subset of the available quantitative and qualitative data is necessarily at best incomplete (and in the worst-case scenario prejudiced, dogmatic, and systematically biased). We are confident that a mature science will sooner or later investigate 5-MeO-DMT in the context of human psychology and physiology. It is just a matter of time — and of neuropolitics… (cf. Rose & Abi-Rached, 2014) We would like to close with an apposite quotation from the distinguished polymath William James who was very interested in mystical/transcendental experiences (as evidenced by his book “The varieties of religious experience”) and who conducted self-experiments with the chemical compound Nitrous Oxide and the psychedelic mescaline containing cactus “Peyote” (*Lophophora williamsii*). James was enthusiastic about the effects of Nitrous Oxide which is not a psychedelic. However, his experiment with Peyote was unfortunately unsuccessful. One can only speculate: Which turn would western psychology have taken if James’ mind would have entered the psychedelic realm? In his classic “Essays in Radical Empiricism” James eloquently articulated the importance of unbiased empirical inquiry:

> “*To be radical, an empiricist must neither admit into his constructions any element that is not directly experienced, nor exclude from them any element that is directly experienced* (James, 1912/1976, p.42).

## Footnotes

1 This present epoch is also termed the 6^th^ mass extinction due to the apid anthropogenic biodiversity loss which is comparable to other exogenously caused mass extinctions in the history of the planet (Régnier et al., 2015; Worm et al., 2006). That is, we are currently witnessing the first mass extinction caused by the behavior of a species. For instance, the last Cretaceous-Paleogene extinction even was with high likelihood caused by the impact of a meteorite or comet.

2 In fact, the term “capitalocene” has been proposed as more accurate (Altvater, 2016). Human pressure on the Earth System is primarily due to the wealthy OECD countries. Their “ecological footprint” (cf. Dietz, Rosa, & York, 2007) is proportionally much larger in comparison to the rest of the world to due consumption (an waste) of resources, i.e., equity significantly factors into the equation of “the great acceleration” (Steffen, Broadgate, Deutsch, Gaffney, & Ludwig, 2015). This might sound like a somewhat extreme political science perspective, but it is important to emphasize that the problem has interdisciplinary dimensions and cannot be fragmented - an interdisciplinary system theoretical approach is needed. The monetary system, economics, philosophy, ethics, and morality play an important role as do the social and political sciences and the neurosciences. A creative solution has to have interdisciplinary ramifications. In fact neoliberalism has been critically discussed in the context of creativity (Gormley, 2018; Harvey, 2007).

3 The term “radical” is etymologically derived from the Latin word “radix” meaning “root” (cf. the radical sign √ in mathematics). That is, radical solution means a “solution which targets the roots of the problem” (the roots are psychological).

4 URL: https://thebulletin.org/2018-doomsday-clock-statement

5 The binomial taxonomical nomenclature (introduced by Carl Linnæus) is etymologically derived from the Latin “homō” meaning “human being” and “sapiēns” meaning “wise” — thus the “wise human”. By contrast, the neologism *Homō consumens* has been proposed as a more accurate/realistic designation given the production and consumption-orientation of the species (Fromm, 1976).

6 Realistic thinkers have argued that the chances of species survival are *de facto* minute (Fromm, 1962). However, classical game theoretical calculi are not applicable to this situation. Even if the chance of success is <1% humanity needs to mobilize all its resources to come up with a solution to the problem of self-destruction.

7 Source: New York Times - May 25 1946, p.13 - “Atomic Education Urged by Einstein” URL: https://nyti.ms/2NpSc8L

8 The etymology of the term is derived from the Ancient Greek ψυχή (psukhḗ, “mind, soul, spirit”) + δῆλος (dêlos, “to manifest, to reveal”), i.e., “psychedelic substances” could be adequately translated as “mind manifesting” or “soul revealing” substances. Similarly decomposed, psychology is “the study of” the “mind, soul, and spirit” — even though most contemporary psychologists would reject this “deep” definition. Previously, psychedelics were also labeled as “psychotomimetics” because they were thought to produce symptoms similar to those of a psychosis. Interestingly, schizophrenia and other psychopathologies involving psychotic symptoms (e.g., bipolar disorder) have been linked to creativity (e.g., Claridge & Blakey, 2009; Power et al., 2015), possibly due to a reduction of latent inhibition (cf. Burch, Hemsley, Pavelis, & Corr, 2006), *inter alia*.

9 It is a plausible hypothesis that psychoactive tryptamines are involved in naturally occurring dream-states. Given its pivotal in biochronological processes, the pineal gland might be an important neuroanatomical locus in this context (cf. Barker, Borjigin, Lomnicka, & Strassman, 2013).

10 It should be emphasized that these chosen examples should not reinforce the superficial conception that creativity only “matters” if it produces material dividends and has no intrinsic value in itself (Fromm, 1976).

11 In a recent randomized double-blind trial ≈70% of participants rated their experimentally induced psychedelic experience as one of their top five spiritually significant lifetime events (Griffiths et al., 2016).

12 An interesting historical fact (especially in the context of ego-dissolution/ego-death) is that Huxley wrote a note to his wife while on his deathbed asking her to inject him with 100μg of LSD (IM). Huxley died while under the influence of the consciousness expanding substance. Another interesting fact is that Huxley was allegedly intimately involved in the illegal CIA MK-ULTRA program which experimented with psychedelic substances on nonconsenting subjects.

13 A connatural conception can also be found in Plato’s “Allegory of the cave” (Republic, 514a-520a). Plato was very much concerned with eternal forms and most mathematicians can be regarded as implicit Platonists (Burnyeat, 2000; Mueller, 2005) even though they might not be explicitly aware of this philosophical heritage (cf. the importance of Δianoia in Plato’s “Theory of Forms” (Cooper, 1966)).

14 The concept of non-duality is closely related the Indian philosophical system of “Advaita Vedānta” (Sanskrit: 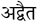 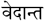, literally, “not-two”) which is one of the most ancient spiritual paths to self-realization. Overcoming/dissolving the illusion of the ego or I-ness principle (Ahaṃkāra) plays a crucial role in this meditative tradition.The experience of ego-dissolution is fundamentally ineffable. Hence, the profundity of ego-dissolution will not be fully comprehended by those readers who have not experiences it first-hand. It relates to the problem of noncommunicable quale: One cannot appreciate the taste of sugar by listening to elaborate descriptions or by studying its molecular structure. One must taste it (cf. Nagel, 1974). In philosophy of mind this is known as the “knowledge argument” (Jackson, 1986).

15 Psychedelic were not only of interest to academic scientists. After initial studies in German concentration camps (e.g., Auschwitz) the CIA developed its own undercover programs (e.g., Project MK-Ultra) in order to test psychedelics compounds on vulnerable and mostly naïve (non-consenting) populations, e.g., prisoners, homeless people, mental patients, etc.

16 John Daniel Ehrlichman who was at this time Assistant to the President (for Domestic Affairs) stated in an interview in 1994 (published in “Harpers” in 2016): *“The Nixon campaign in 1968, and the Nixon White House after that, had two enemies: the antiwar left and black people. You understand what I’m saying? We knew we couldn’t make it illegal to be either against the war or black, but by getting the public to associate the hippies with marijuana and blacks with heroin, and then criminalizing both heavily, we could disrupt those communities. We could arrest their leaders, raid their homes, break up their meetings, and vilify them night after night on the evening news. Did we know we were lying about the drugs? Of course we did.”*

17 In the United Kingdom, the recently ratified “Psychoactive substances act” which reached Royal Assent in January 2016 complicates the matter by creating societal, political, and fiscal impediments to scientific research into the neurobiology of psychedelics. For more information, see: http://www.legislation.gov.uk/ukpga/2016/2/contents/enacted

18 Supra-physiological describes a level of efficacy which is unseen in organisms which evolved according to the principles of natural evolution.

19 Even though the chemical structure of both compounds is very similar their psychological effects are incommensurable.

20 Albert Hofmann (1906-2008) also discovered LSD in 1938 but he was unaware of its psychoactivity until 1943 when he conducted the first self-experiment. Hofmann, who later served as a member of the Nobel Prize Committee, stated on his 100th birthday: “*It gave me an inner joy, an open mindedness, a gratefulness, open eyes and an internal sensitivity for the miracles of creation.* […] *I think that in human evolution it has never been as necessary to have this substance LSD. It is just a tool to turn us into what we are supposed to be*.”

22 For instance, the Pearson correlation coefficient for “global creativity” and OTE is r = .655 and for “creative achievement” r = .481, By contrast, “Math-science creativity” is not statistically significantly correlated with OTE (r =.059; ns; for further correlation between various facets of creativity and the Big Five factors see Silvia, Nusbaum, Berg, Martin, & O’Connor, 2009). The salient correlation between OTE and creativity has been reported in many studies (a pertinent meta-analysis has been conducted by Feist, 1998; a recent study reporting a strong relationship between OTE and creativity has been conducted by Puryear, Kettler, & Rinn, 2017). Furthermore, a meta-analytical structural equation model of 25 independent studies showed that OTE is the strongest FFM predictor of creative self-beliefs (r = .467; Karwowski & Lebuda, 2016).

23 Bertrand Russel discussed the links between mysticism, creative intuition/insight, and logic in great detail in his excellent essay “Mysticism and logic” (Russell, 1981).

24 The rich-club coefficient Φ is a networks metric which quantifies the degree to which well-connected nodes (beyond a certain richness metric) also connect to each other. Hence, the rich-club coefficient can be regarded as a notation which quantifies associativity. Conceptually related research concluded that “associative abilities represent valid elementary cognitive abilities underlying creativity” (Benedek, Könen, & Neubauer, 2012).

25 Furthermore, the authors argue convincingly that the notion that LSD (and other psychedelics) “expand” consciousness is quantitatively supported by their data. Specifically, they argue that the neurophysiological changes associated with psychedelic states contrast with states of diminished consciousness (e.g., deep sleep or general anesthesia). The obtained results are congruent with the idea that psychedelic and unconscious states can be conceptualized as polar-opposites on a continuous spectrum of conscious states. Furthermore, the authors suggest that the level of consciousness is quantitatively determined by the level of neuronal entropy (in accord with the entropic brain hypothesis formulated by Carhart-Harris et al., 2014). It has been suggested that Aldous Huxley “reduction valve” hypothesis appears to be relevant in this context.

26 Recent evidence focusing on changes in the coupling of electrophysiological brain oscillations by means of transfer entropy suggests that serotonergic psychedelics temporarily change information transfer (via an increase of entropy?) within neural hierarchies by decreasing frontal of top-down control, thereby releasing posterior bottom-up information transfer from inhibition (Francesc Alonso, Romero, Angel Mañanas, & Riba, 2015).

27 A possible neural mechanism might be found in the entropic brain hypothesis (Carhart-Harris et al., 2014; Lebedev et al., 2016). Pertinent experimental evidence comes from a recent magnetoencephalographic (MEG) study which showed that classical psychedelics increase signal diversity (Schartner, Carhart-Harris, Barrett, Seth, & Muthukumaraswamy, 2017), a quantitative finding which appears highly relevant in the context of contemporary creativity research.

28 Interestingly, ego-dissolution was also statistically significantly correlated with enhanced well-being/life-satisfaction (ρ = 0.392). For alcohol (ρ = −0.112) and cocaine (ρ = −0.083) this positive effect was absent. Due to the quasi-experimental nature of this study no solid conclusions are possible. Systematic experimental research is needed to elucidate this important topic which has obvious societal relevance.

29 The snuff was administered in a ceremonial setting in which the ground seeds of the cojóbana tree (*Anadenanthera peregrina*) were inhaled via a Y-shaped pipe called Cohoba (Ortiz, 1941).

30 For example, toad-effigies and iconography (with accentuated glands) are found in archaeological excavation from ancient Mayan and Aztec cultures, e.g., artworks of “Tlaltecuhtli” (Aztec: 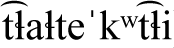) - the earth or earth mother as a monstrous toad (Furst, 1972).

31 See, for example, https://www.fractalimagination.com Interestingly, under the influence of low doses of LSD spiders spin webs of greater regularity (Witt, 1951). Other researchers applied fractal theory to investigate “the correlation between the fractal structure of spider’s web and the fractal dynamics of its brain signal” (Namazi, 2017).

32 The long history of human usage of this naturally occurring compound in various cultures suggests that it does not convey a significant disadvantage in terms of evolutionary fitness i.e., mutation/natural selection (cf. Martin & Nichols, 2017). Profit-oriented pharmaceutical companies, on the other hand, actively market patented synthetic designer drugs which do not have any evolutionary track record and might cause all kinds of unforeseen neurological, genetic, and epigenetic problems in the long run (cf. Kim et al., 2009), for instance, the widespread prescription of methylphenidate (e.g., Ritalin) in preschool children (Keane, 2008), based on questionable DSM-5 nosology (Phillips et al., 2012b, 2012c, 2012d, 2012a). In contrast to patentable psychopharmacological agents, there is no revenue model for naturally occuring psychedelics in the merely profit-oriented capitalistic paradigm.

33 An animal neuroimaging study conducted by Riga et al. (2014) showed that 5-MeO-DMT decreased BOLD responses in the striate cortex (V1) and the medial prefrontal cortex (mPFC).

34 Note that this statement is not an objective empirically validated ontological fact. It is based on qualitative phenomenological experiences often induced by ego-dissolution (e.g., caused by meditation, introspection, psychedelics, spontaneous epiphany, etc.). Ego-less pure awareness plays a central role in many ancient philosophical schools of thought (Mahayana and Zen Buddhism, Advaita Vedanta, Taoism, Sufism, ect. pp.).

35 Indeed, anecdotal reports indicate that it can cause glossolalia (in a addition to synaesthesia and other remarkable effects which are of importance for creativity research).

36 It has been argued elsewhere that “increased creativity may […] constitute a manifestation of posttraumatic growth, defined as retrospective perceptions of positive psychological changes that take place following experiences of highly challenging life circumstances” (Forgeard, 2013, p. 245).

37 Interestingly, preliminary evidence suggests that it is effective in the treatment of addiction, depression, and obsessive-compulsive disorders (Bogenschutz et al., 2015; Carhart-Harris et al., 2016). This is congruent with the formulated idea that 5-MeO-DMT has the potential to change persistent habitual modes of thought.

38 This idea could be empirically tested, for instance, by utilizing a semantic priming paradigm in order to investigate spread of activation (as proxy for verbal creativity). Exemplary studies have been conducted with the dopamine precursor L-Dopa by, for example, Kischka et al. (1996) in order to elucidate the role of dopaminergic neurotransmission in verbal creativity. Anecdotal evidence suggest that serotonergic psychedelics can enhance verbal creativity significantly (longitudinally). In the acute phase, many psychedelics interfere strongly with the linguistic system (a breakdown of semantic and syntactic facilities is oftentimes reported). Interesting, glossolalia is reported in a few cases.

39 It should be noted that psychedelic might cause serious psychological harm to certain populations with psychopathological dispositions (e.g., specific 5-HT receptor polymorphism). Ergo, careful a priori screening is crucial for ethically responsible research.

40 The concept of interconnectedness is of utmost importance from an ecopsychology point of view (cf. Key & Kerr, 2011). The formulated hypothesis thus has significant real-world societal significance. The illusion of disconnection from nature (Fromm, 1962) lies at the root of many destructive human behaviors which have far reaching detrimental consequences (individual and society, micro and macro are not separable - therefore individual changes translate into global changes). Impetus for the hypothesis at hand is partially derived from recent studies which indicate that classical psychedelics increase nature-relatedness (Forstmann & Sagioglou, 2017; Lyons & Carhart-Harris, 2018b).

41 Alcohol, which is legal and indeed systematically promoted by the alcohol industry, has a very unsafe LD_50_ profile and is proven to be neurotoxic (Da Lee et al., 2005; Jacobus & Tapert, 2013). Recent longitudinal research has shown that even moderate alcohol consumption has detrimental effects on various neuroanatomical structures (e.g., hippocampal atrophy). Psilocybin, on the other hand, has been shown to induce neurogenesis in the hippocampus in animal studies (Catlow, Song, Paredes, Kirstein, & Sanchez-Ramos, 2013).

42 For more information see http://www.legislation.gov.uk/ukpga/2016/2/contents/enacted

43 In an age in which public opinions are systematically manipulated (Bernays, 1928; L’Etang, 1999) and “consent is manufactured” (Chomsky, 1992; Fleming & Oswick, 2014) cognitive diversity is a disruptive factor which might interfere with the smooth workings of the “mega-machine” (cf. Fromm, 1962; Mumford, 1967).

44 For more information on the declassified “U.S. torture manuals” see https://nsarchive2.gwu.edu//nsa/archive/news/dodmans.htm

45 See letter by the former APA President Alan E. Kazdin to George Bush: https://www.apa.org/news/press/releases/2008/10/bush-interrogations.aspx

46 See official statement of the Bulletin of Atomic Scientists. URL: https://thebulletin.org/2018-doomsday-clock-statement

47 As Bertrand Russel put is: “The whole duality of mind and matter […] is a mistake; there is only one kind of stuff out of which the world is made, and this stuff is called mental in one arrangement, physical in the other.” (Russell, 1913, p.15). Russel’s monism stands in sharp contrast with the (mainly unquestioned) “reductive materialism” working-hypothesis which forms the predominant basis of contemporary science.

48 This view has been deeply challenged by contemporary experimental quantum physics (Handsteiner et al., 2017; Hensen et al., 2015).

49 This view is gradually changing, for instance, Christof Koch stated in a 2014 SCIENTIFIC AMERICA article that “the mental is too radically different for it to arise gradually from the physical” (p. 2).

50 Cf. The widely studied status quo bias (Fleming, Thomas, & Dolan, 2010)

